## Supplementary information for "Structural insights into complex I deficiency and assembly from the disease-related *ndufs4*^-/-^ mouse"

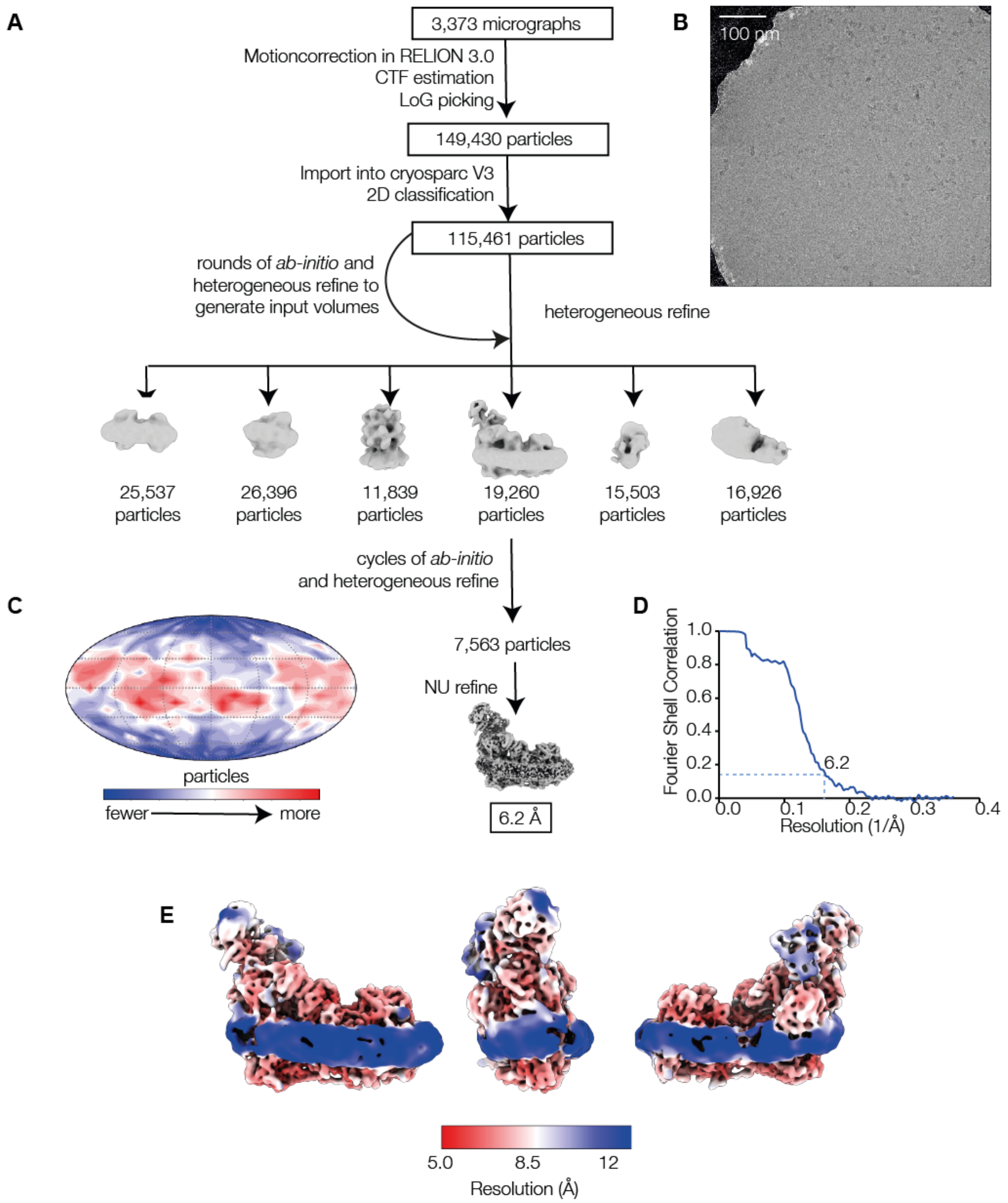


**Figure S1. Cryo-EM data collection and structural reconstruction of complex I from *ndufs4*$^{-/-}$ mouse kidney.** (A) Data processing scheme for the mouse complex I from *ndufs4*$^{-/-}$ mouse kidney; B) example micrograph; C) angular distribution of particles; D) the global resolution estimate from the masked Fourier Shell Correlation curve is 6.2 Å at FSC = 0.143; and E) local resolution map calculated in cryoSPARC using an FSC = 0.5 cut-off.

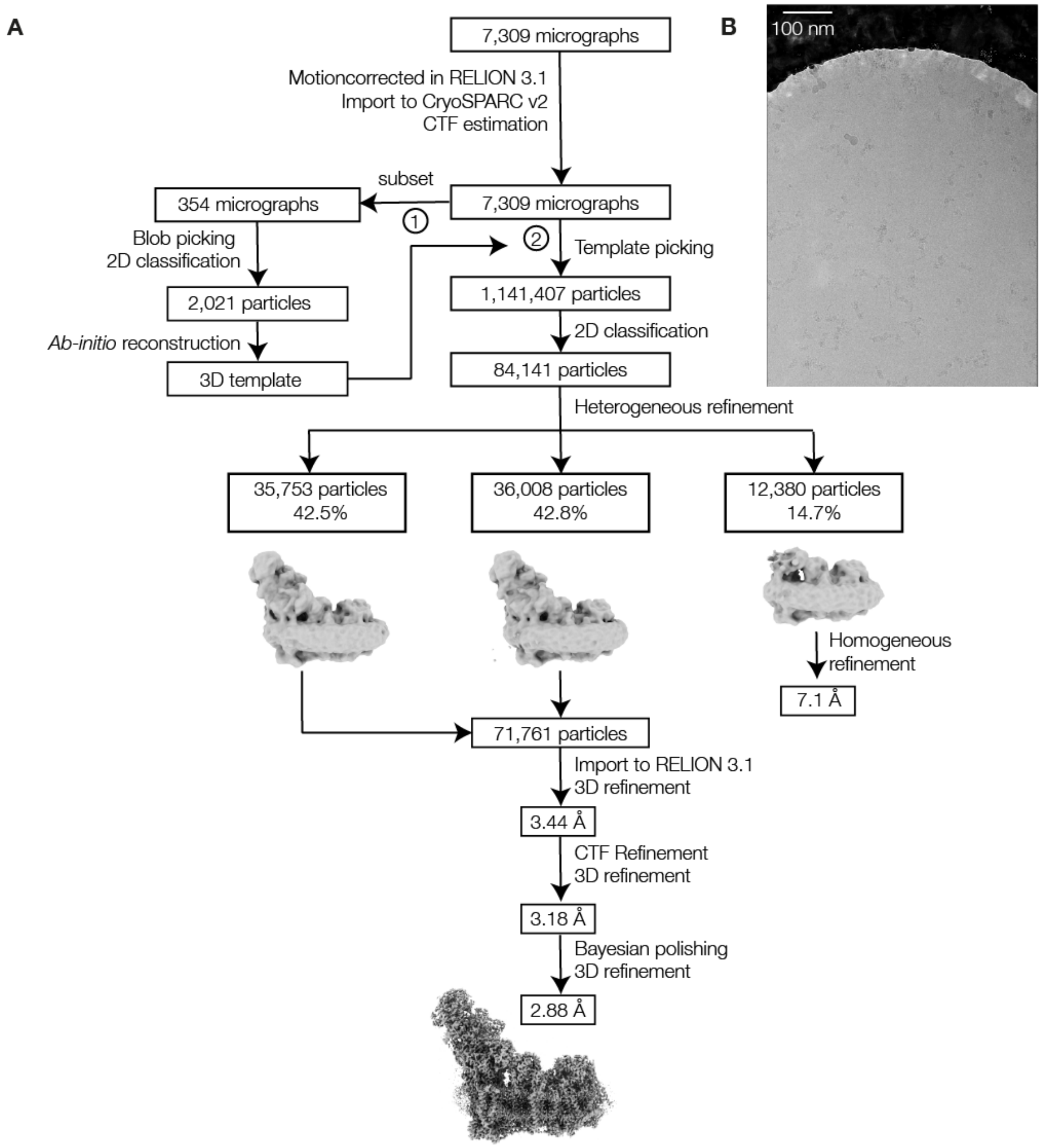


**Figure S2. Cryo-EM data collection and structural reconstruction of complex I from *ndufs4$^{-/-}$* mouse heart.**

A) Data processing scheme for the mouse complex I from *ndufs4$^{-/-}$* mouse heart; B) example micrograph.

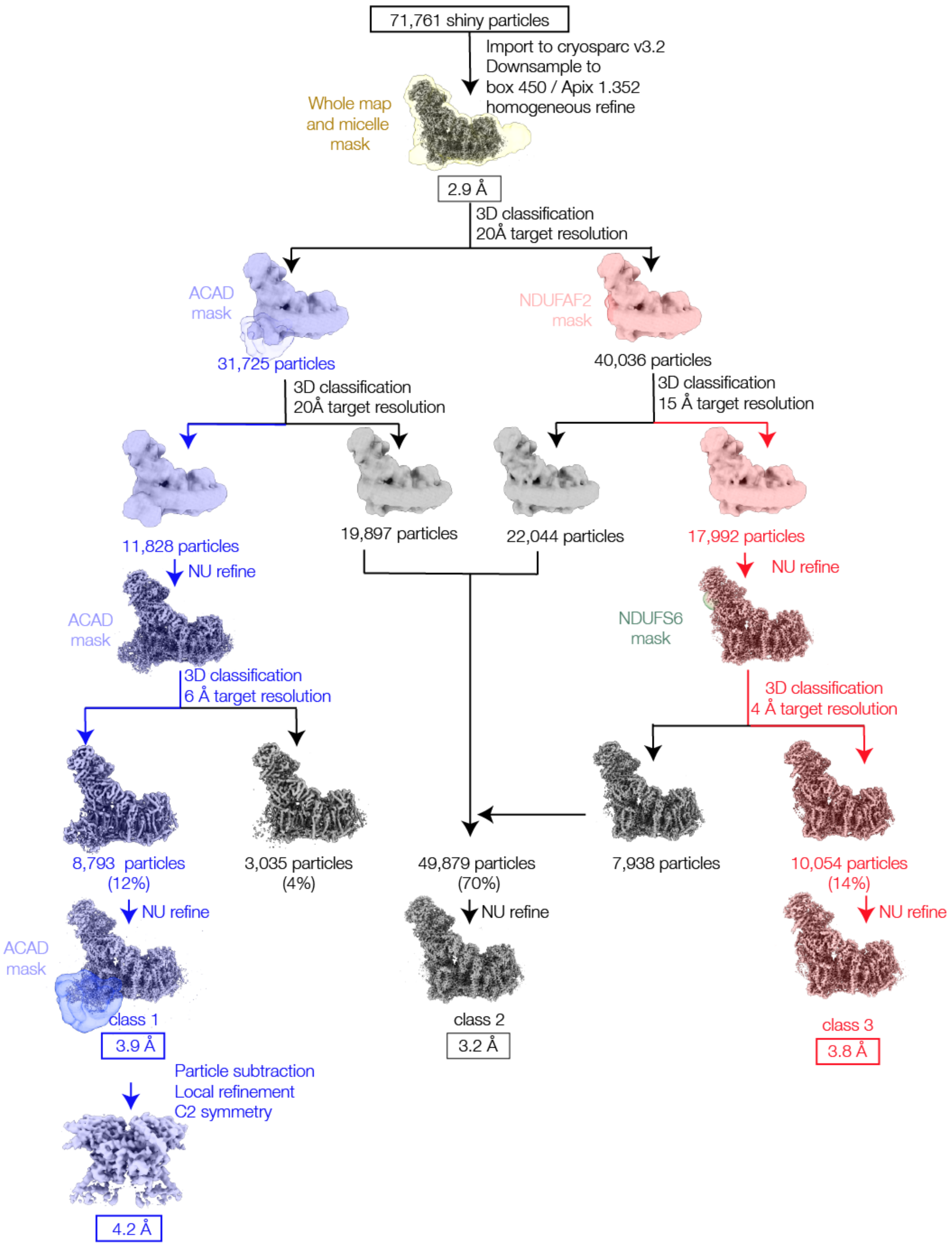


**Figure S3. Subclassification of polished particles in cryoSPARC by 3D classification**. After two rounds of classification (blue and pink) the intermediate (top) pink species had poor density for both NDUFS6 and NDUFAF2. Further subclassification around subunit NDUFS6 then showed that the molecules in the pink population contain either NDUFS6 (grey) or NDUFAF2 (pink), but not both.

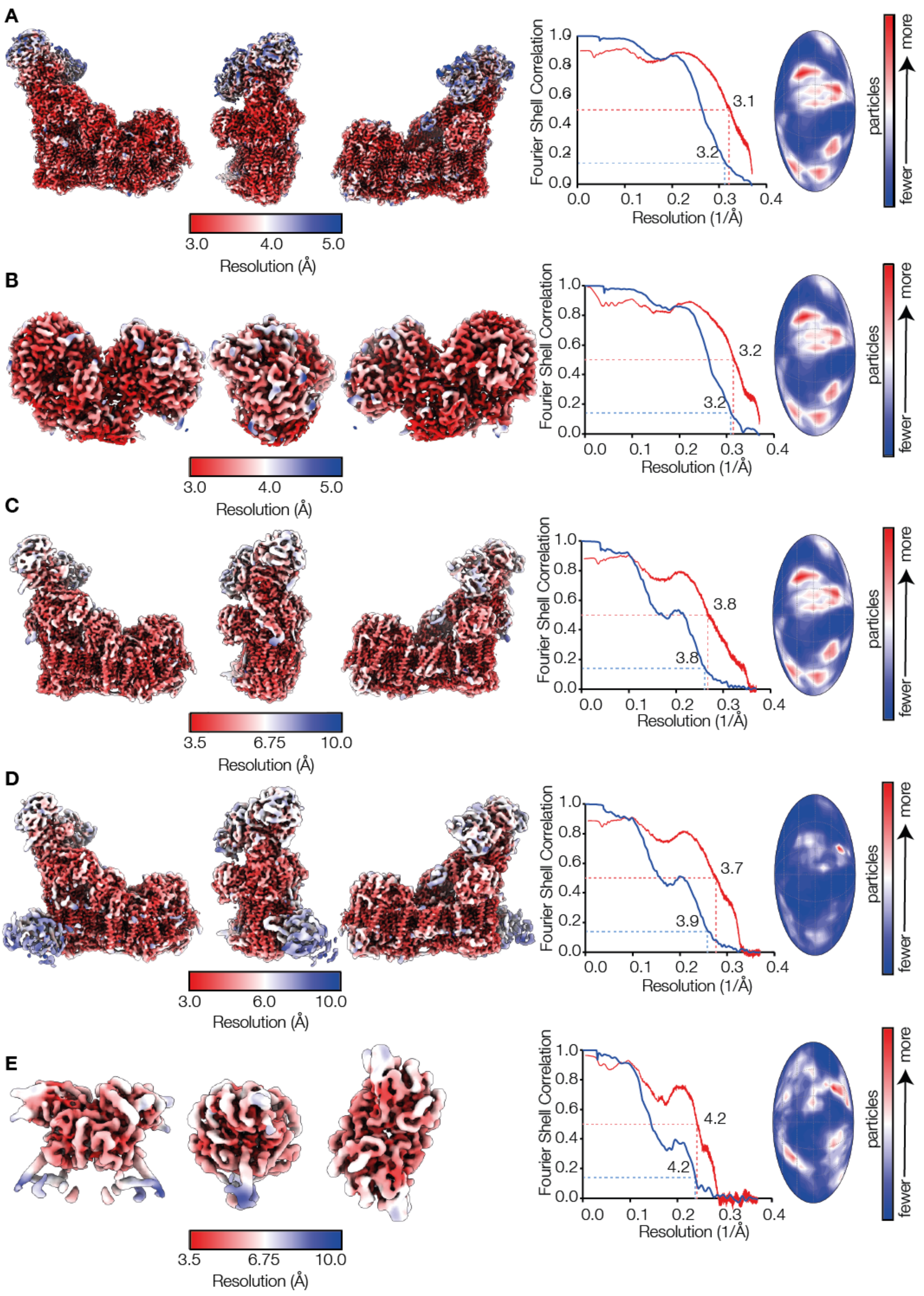


**Figure S4. Local resolution maps, FSC curves and angular distribution plots for structural data on *ndufs4*$^{-/-}$ heart complex I.** Local resolutions were calculated in cryoSPARC using an FSC = 0.5 cut-off. Half-map (blue) and map-model (red) FSC curves are shown with the resolution at FSC = 0.143 indicated for the half map FSC, and FSC = 0.5 is indicated for the map-model FSC. Data are shown for A) class 2 *ndufs4*$^{-/-}$ complex I; B) class 2 *ndufs4*$^{-/-}$ N-module of complex I; C) class 3 *ndufs4*$^{-/-}$ complex I; D) class1 *ndufs4*$^{-/-}$ complex I; E) class 1 *ndufs4*$^{-/-}$ ACADVL only.

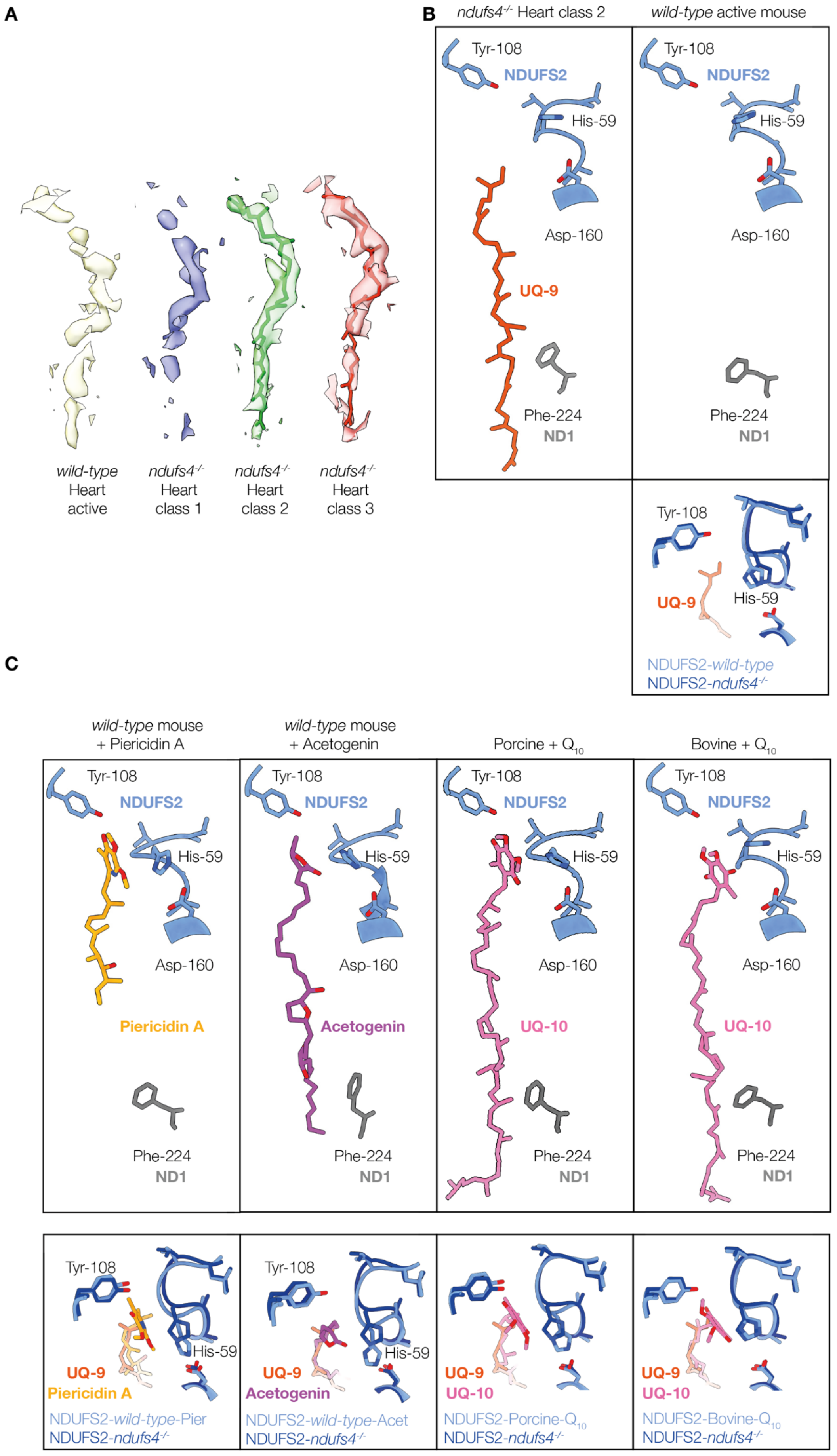
A
wild-type Heart active
ndufs4-/- Heart class 1
ndufs4-/- Heart class 2
ndufs4-/- Heart class 3
B
ndufs4-/- Heart class 2
wild-type active mouse
Tyr-108
NDUFS2
His-59
Asp-160
UQ-9
Phe-224
ND1
NDUFS2-wild-type
NDUFS2-ndufs4-/-
C
wild-type mouse + Piericidin A
wild-type mouse + Acetogenin
Porcine + Q10
Bovine + Q10
Piericidin A
Acetogenin
UQ-10
NDUFS2-wild-type-Pier
NDUFS2-wild-type-Acet
NDUFS2-Porcine-Q10
NDUFS2-Bovine-Q10

**Figure S5. Ubiquinone binding in the *ndufs4*$^{-/-}$ heart complex I.** A) Cryo-EM density in the ubiquinone-binding channel in the density maps for *wild-type* active mouse complex I (EMD: 11377 (Bridges *et al*, 2020)) and *ndufs4*$^{-/-}$ heart complex I classes 1 to 3 (EMD: 16398, 16516, 16518). B) Key ubiquinone-binding site elements in *ndufs4*$^{-/-}$ heart class 2 (PDB: 8CA3) and the active *wild-type* reference structure (PDB: 6ZR2) (Bridges *et al*, 2020) and a comparison of structures in the headgroup-binding region. C) Key ubiquinone-binding site elements in structures of *wild-type* mouse with piericidin (PDB: 6ZTQ) (Bridges *et al*, 2020), *wild-type* mouse with acetogenin (PDB: 7PSA) (Grba *et al*, 2022), bovine complex I with ubiquinone-10 (7QSL) (Chung *et al*, 2022b), and porcine complex I with ubiquinone-10 (PDB: 7V2C) (Gu *et al*, 2022), and comparisons of structures with *ndufs4*$^{-/-}$ heart class 2 in the headgroup-binding region.

| Subunit | MALDI TOF/TOF MS/MS | | | LC MS | | |
|---|---|---|---|---|---|---|
| | Unique peptides | Sequence coverage | Score | Unique peptides | Sequence coverage | Score/32 |
| **NDUFA1** | 4 | 42% | 76/34 | 5 | 42% | 109 |
| **NDUFA2** | 4 | 32% | 238/33 | 9 | 70% | 161 |
| **NDUFA3** | 3 | 32% | 40/34 | 4 | 50% | 74 |
| **NDUFA5** | 3 | 31% | 109/35 | 10 | 63% | 318 |
| **NDUFA6** | 3 | 26% | 148/33 | 19 | 87% | 546 |
| **NDUFA7** | - | - | - | 18 | 85% | 613 |
| **NDUFA8** | 4 | 26% | 127/34 | 11 | 48% | 333 |
| **NDUFA9** | 11 | 35% | 596/35 | 25 | 64% | 1050 |
| **NDUFA10** | 11 | 42% | 627/34 | 20 | 41% | 699 |
| **NDUFA11** | - | - | - | 2 | 10% | 146 |
| **NDUFA12** | - | - | - | 13 | 77% | 460 |
| **NDUFA13** | 5 | 30% | 230/35 | 20 | 81% | 1165 |
| **NDUFAB1** | - | - | - | 1 | 11% | 55 |
| **NDUFB1** | 3 | 45% | 116/34 | 5 | 50% | 56 |
| **NDUFB2** | 2 | 8% | 70/33 | 1 | 8% | 41 |
| **NDUFB3** | 2 | 14% | 39/35 | 5 | 33% | 88 |
| **NDUFB4** | 7 | 53% | 302/33 | 17 | 69% | 488 |
| **NDUFB5** | - | - | - | 9 | 35% | 316 |
| **NDUFB6** | - | - | - | 15 | 75% | 260 |
| **NDUFB7** | - | - | - | 5 | 45% | 405 |
| **NDUFB8** | 5 | 38% | 273/34 | 9 | 43% | 479 |
| **NDUFB9** | 3 | 24% | 78/34 | 16 | 66% | 381 |
| **NDUFB10** | 1 | 8% | 54/34 | 11 | 55% | 400 |
| **NDUFB11** | 1 | 7% | 41/30 | 11 | 56% | 472 |
| **NDUFC2** | 3 | 17% | 126/33 | 12 | 60% | 251 |
| **NDUFS1** | 6 | 13% | 245/32 | 44 | 47% | 2231 |
| **NDUFS2** | 12 | 30% | 476/35 | 23 | 48% | 809 |
| **NDUFS3** | 6 | 30% | 336/34 | 18 | 46% | 796 |
| **NDUFS5** | - | - | - | 10 | 46% | 195 |
| **NDUFS6** | 4 | 39% | 189/35 | 6 | 50% | 466 |
| **NDUFS7** | 7 | 14% | 219/34 | 10 | 30% | 332 |
| **NDUFS8** | - | - | - | 8 | 30% | 173 |
| **NDUFV1** | 11 | 27% | 475/35 | 24 | 41% | 731 |
| **NDUFV2** | - | - | - | 13 | 45% | 804 |
| **NDUFV3** | - | - | - | 6 | 45% | 121 |
| **ND1** | - | - | - | 3 | 8% | 116 |
| **ND2** | 1 | 4% | 61/33 | - | - | - |
| **ND4** | 4 | 7% | 112/35 | 3 | 7% | 73 |
| **ND5** | 1 | 2% | 50/34 | 5 | 8% | 217 |
| **ACAD9** | 2 | 4% | 37/33 | 4 | 8% | 147 |
| **ACADVL** | 10 | 21% | 407/34 | 28 | 45% | 1241 |
| **NDUFAF2** | 3 | 23% | 149/35 | 11 | 71% | 273 |

**Table S1. Peptide-based protein identification of the composition of *ndufs4*$^{-/-}$ complex I purified from heart.** The number of peptides detected from each subunit is given together with the sequence coverage, relative to the sequence of the immature protein. The score given for each subunit is the sum of the peptide scores, with the denominator representing the 95% confidence threshold ($p < 0.05$). Subunits NDUFC1, ND3, ND4L and ND6 (as well as NUDFS4) were not detected.

| Subunit | MALDI TOF/TOF MS/MS | | | LC MS | | |
|---|---|---|---|---|---|---|
| | Unique peptides | Sequence coverage | Score | Unique peptides | Sequence coverage | Score/21 |
| **NDUFA1** | 2 | 14% | 57/34 | 3 | 32% | 63 |
| **NDUFA2** | 5 | 38% | 191/33 | 10 | 72% | 351 |
| **NDUFA3** | 2 | 21% | 74/34 | 1 | 10% | 26 |
| **NDUFA5** | 3 | 45% | 209/34 | 3 | 29% | 100 |
| **NDUFA6** | 4 | 32% | 203/32 | 12 | 80% | 520 |
| **NDUFA7** | - | - | - | 13 | 78% | 725 |
| **NDUFA8** | 5 | 39% | 275/33 | 3 | 16% | 136 |
| **NDUFA9** | 13 | 37% | 836/34 | 15 | 51% | 608 |
| **NDUFA10** | 11 | 40% | 723/33 | 6 | 21% | 237 |
| **NDUFA11** | - | - | - | - | - | - |
| **NDUFA12** | 2 | 6% | 59/33 | 12 | 64% | 400 |
| **NDUFA13** | 3 | 22% | 138/35 | 13 | 71% | 649 |
| **NDUFAB1** | 4 | 21% | 216/34 | - | - | - |
| **NDUFB1** | 2 | 31% | 85/33 | 3 | 33% | 58 |
| **(NDUFB2)** | 2 | 8% | 52/33 | 1 | 8% | 61 |
| **(NDUFB3)** | 3 | 14% | 50/33 | 4 | 22% | 74 |
| **NDUFB4** | 6 | 46% | 298/33 | 9 | 61% | 427 |
| **NDUFB5** | 3 | 19% | 114/33 | 6 | 32% | 175 |
| **NDUFB6** | 5 | 40% | 175/33 | 9 | 57% | 292 |
| **NDUFB7** | 2 | 10% | 134/30 | 2 | 25% | 227 |
| **NDUFB8** | 8 | 49% | 449/33 | 5 | 37% | 302 |
| **NDUFB9** | - | - | - | 9 | 50% | 220 |
| **NDUFB10** | 4 | 21% | 138/33 | 7 | 40% | 271 |
| **NDUFB11** | - | - | - | 7 | 42% | 165 |
| **NDUFC2** | 4 | 27% | 166/34 | 7 | 45% | 340 |
| **NDUFS1** | 12 | 20% | 653/32 | 32 | 41% | 1653 |
| **NDUFS2** | 10 | 31% | 729/33 | 14 | 34% | 519 |
| **NDUFS3** | 12 | 49% | 818/33 | 10 | 33% | 509 |
| **NDUFS4** | 7 | 30% | 449/33 | 8 | 34% | 284 |
| **NDUFS5** | 1 | 9% | 85/34 | 5 | 43% | 73 |
| **NDUFS6** | 4 | 32% | 278/33 | 4 | 41% | 258 |
| **NDUFS7** | 6 | 14% | 217/33 | 5 | 24% | 280 |
| **NDUFS8** | 5 | 24% | 318/32 | 3 | 16% | 136 |
| **NDUFV1** | 12 | 27% | 574/33 | 18 | 32% | 445 |
| **NDUFV2** | 7 | 25% | 311/33 | 8 | 33% | 526 |
| **NDUFV3** | - | - | - | 3 | 25% | 96 |
| **ND4** | 4 | 7% | 149/33 | - | - | - |
| **ND5** | 3 | 7% | 84/32 | - | - | - |
| **ACADVL** | 12 | 25% | 796/33 | 16 | 30% | 777 |
| **(NDUFAF2)** | 4 | 22% | 80/33 | | | |

**Table S2. Peptide-based protein identification of the composition of *wild-type* complex I purified from heart.** The number of peptides detected from each subunit is given together with the sequence coverage, relative to the sequence of the immature protein. The score given for each subunit is the sum of the peptide scores, with the denominator representing the 95% confidence threshold ($p < 0.05$). For proteins labelled in brackets individual peptide scores were below the 95% threshold but their protein scores were above the peptide threshold. Subunits NDUFC1, ND1, ND2, ND3, ND4L and ND6 (as well as ACAD9) were not detected.

| | *Ndufs4* $^{-/-}$ kidney | *Ndufs4* $^{-/-}$ heart | | | | |
|---|---|---|---|---|---|---|
| | | **Class 1** | **Class 1 ACADVL** | **Class 2** | **Class 2 N-module** | **Class 3** |
| PDB code | - | 8C2S | 8CA1 | 8CA3 | 8CA4 | 8CA5 |
| EMDB code | 16514 | 16398 | 16515 | 16516 | 16517 | 16518 |
| **Data collection and processing** | | | | | | |
| Magnification (nominal) | 59,000 | 64,000 | | | | |
| Voltage (kV) | 300 | 300 | | | | |
| Electron exposure ($e^{-}/Å^2$) | 49.24 | 45 | | | | |
| Defocus range (μm) | -2.1 to -3.3 | -1.5 to -2.9 | | | | |
| Calibrated Pixel size (Å) | 1.39 | 1.352 | | | | |
| Symmetry imposed | C1 | C1 | C2 | C1 | C1 | C1 |
| Initial particle images | 149,198 | 1,141,407 | | | | |
| Final particle images | 7,563 | 8,793 | 8,793 | 50,914 | 50,914 | 10,005 |
| Map sharpening B-factor ($Å^2$) | -130 | -10 | -27 | -27 | -10 | 0 |
| Map resolution (FSC = 0.143) (Å) | 6.2 | 3.9 | 4.3 | 3.2 | 3.2 | 3.9 |
| **Model Refinement** | | | | | | |
| Initial model used | | 6ZR2 | AF-P50544 | 6ZR2 | 6ZR2 | 6ZR2<br>AF-Q59J78 |
| Model resolution (FSC = 0.5) (Å) | | 3.7 | 4.2 | 3.1 | 3.2 | 3.7 |
| Model composition | | | | | | |
| Non-hydrogen atoms | | 63,713 | 9,036 | 63,713 | 11,113 | 64,198 |
| Protein residues | | 7,799 | 1,178 | 7,799 | 1,439 | 7,851 |
| Ligands | | 32 | 2 | 32 | 6 | 31 |
| B factors mean ($Å^2$) | | | | | | |
| Protein | | 62.57 | 102.07 | 51.34 | 56.77 | 76.25 |
| Ligand | | 63.68 | 98.64 | 60.08 | 5.39 | 77.85 |
| RMSD Bond lengths (Å) | | 0.007 | 0.006 | 0.005 | 0.005 | 0.004 |
| RMSD Bond angles (°) | | 1.071 | 1.185 | 0.995 | 1.017 | 0.658 |
| Validation | | | | | | |
| MolProbity score | | 1.94 | 1.89 | 1.68 | 1.87 | 1.90 |
| Clashscore | | 10.15 | 9.75 | 6.14 | 7.44 | 10.21 |
| Poor rotamers (%) | | 0.16 | 0 | 0.06 | 0.25 | 0.25 |
| EMRinger score | | 2.15 | 2.19 | 3.03 | 3.01 | 2.19 |
| Ramachandran plot | | | | | | |
| Favored (%) | | 93.80 | 94.38 | 95.12 | 92.65 | 94.54 |
| Allowed (%) | | 6.07 | 5.45 | 4.85 | 7.28 | 5.34 |
| Disallowed (%) | | 0.13 | 0.17 | 0.04 | 0.07 | 0.12 |
| Z-score (whole) | | 0.35 | 1.92 | 0.98 | 0.34 | 0.84 |
| Map-model correlation $CC_{mask}$ | | 0.82 | 0.82 | 0.87 | 0.89 | 0.78 |

**Table S3. Cryo-EM data collection, refinement and validation statistics for maps and models of *ndufs4*$^{-/-}$ complex I and associated proteins.**

| | ***Wild-type* active**<br>EMD-11377 | ***Wild-type* deactive**<br>EMD-11810 |
|---|---|---|
| ***ndufs4$^{-/-}$* class 1**<br>EMD-16398 | 94 | 81-83 |
| ***ndufs4$^{-/-}$* class 2**<br>EMD-16516 | 95-97 | 81-85 |
| ***ndufs4$^{-/-}$* class 3**<br>EMD-16518 | 94-95 | 87-88 |

**Table S4. Map-map correlations for consensus *ndufs4$^{-/-}$* class 1-3 maps against reference *wild-type* active and deactive maps.** All maps were first lowpass-filtered to 3.9 Å. Maps thresholds were class 1, 2.07; class 2, 1.83; class 3, 1.85; *wild-type* active, 0.022; *wild-type* deactive, 0.04.
